# BarQC: Quality Control and Preprocessing for SPLiT-Seq Data

**DOI:** 10.1101/2025.02.04.635005

**Authors:** Maria Rossello, Sophie Tandonnet, Isabel Almudi

**Affiliations:** Department of Genetics, Microbiology and Statistics, Universitat de Barcelona, Diagonal 643, 08028 Barcelona, Spain; Institut de Recerca de la Biodiversitat (IRBio), Universitat de Barcelona, Diagonal 643, 08028 Barcelona, Spain

## Abstract

**Summary:** SPLiT-seq is a cost-effective single-cell RNA-sequencing method, easy to implement in most laboratories. However, the data preprocessing is not straightforward, and no solution currently exists to perform a basic quality control of the data. Here, we present BarQC, a quality control tool, which identifies and corrects barcodes in SPLiT-seq data and provides a report on the distribution of each barcode and UMI as well as an estimation of the number of cells present in the data. Additionally, BarQC can be incorporated in the preprocessing pipeline by constructing a sorted BAM file tagged with each barcode sequence.

**Availability of Implementation:** BarQC was implemented in Python3, and its source code is freely available on GitHub at https://github.com/mayflylab/barQC/ under the MIT Licence. BarQC has been tested on a Linux server.

## Introduction

The recent explosion of single-cell sequencing studies, possible thanks to technological advancements and dropping sequencing prices, is paving the way towards a new, cell-level understanding of biological processes. It is now possible to build comprehensive catalogues of cell types (Plass *et al*., 2018; Sebé-Pedrós *et al*., 2018; Li *et al*., 2022; Konstantinides *et al*., 2022; Aldridge and Teichmann, 2020), to infer links between them, to identify common or unique processes and gene modules among groups of cells (Najle *et al*., 2023; Álvarez-Campos *et al*., 2024), and to compare cell compositions between species, sexes, conditions, tissue type, developmental stages, or cellular state (Li *et al*., 2024; Salamanca-Díaz *et al*., 2024; Hao *et al*., 2024; Elmentaite *et al*., 2022). Single-cell approaches also allow us to identify new molecular actors and processes and characterize new populations or subpopulations of cells (Jindal *et al*., 2018; Drayman *et al*., 2019; Konstantinides *et al*., 2022). The applications of single-cell methods range from fundamental biological questions to understanding diseases (e.g. cancer cells (Tirosh and Suva, 2024)) as well as assessing environmental health and function (e.g. microbial communities Kuchina *et al*., 2021).

Over the last decade, a number of methods have arisen to obtain sequencing at the cell level, including drop-seq (Macosko *et al*., 2015), SMART-seq (Picelli *et al*., 2014; Ramsköld *et al*., 2012), and SPLiT-seq (Rosenberg *et al*., 2018). In this note, we will focus on SPLiT-seq, a method offering numerous advantages. SPLiT-seq requires no special equipment, enables the sequencing of hundreds of thousands of cells simultaneously via Illumina paired-end sequencing, and is cost-effective, with the price per cell decreasing as the number of cells increases. At its core, SPLiT-seq uses the cell itself as the compartment isolating RNA molecules from other cells (Rosenberg *et al*., 2018). Briefly, the method relies on uniquely tagging the transcripts of each cell through the sequential addition of barcodes (short DNA sequences) (See Figure 1A for a diagram of the library structure). Each unique barcode combination is characteristic of a single cell, allowing multiplexing of various samples and conditions within a single experiment. Low collision rates are achieved by limiting the number of cells per well and leveraging the exponential number of barcode combinations created at each pooling/splitting step (Rosenberg *et al*., 2018). As all samples of an experiment can be processed at the same time, the SPLiT-seq method is less prone to batch effects. Despite its advantages making it an accessible and scalable solution for researchers, the processing of SPLiT-seq reads -- which is a critical step to build the matrix of expression -- lacks an easy-to-use pipeline combined with a comprehensive Quality Control (QC) of the data. Most methods available, such as STARsolo (Dobin *et al*., 2013), split-seq-pipeline (Github: yjzhang/split-seq-pipeline), zUMIs (Parekh *et al*., 2018) and alevin-fry splitp (Github: COMBINE-lab/splitp), are strict in identifying the barcodes as they usually rely on the barcodes’ theoretical known positions in the read (Fixed position). This approach lacks the flexibility and sensitivity to recover reads affected by small insertions or deletions (indels) that can shift barcode starting positions. Moreover, these methods do not offer a visual overview of the SPLiT-seq data.

**Figure 1.**
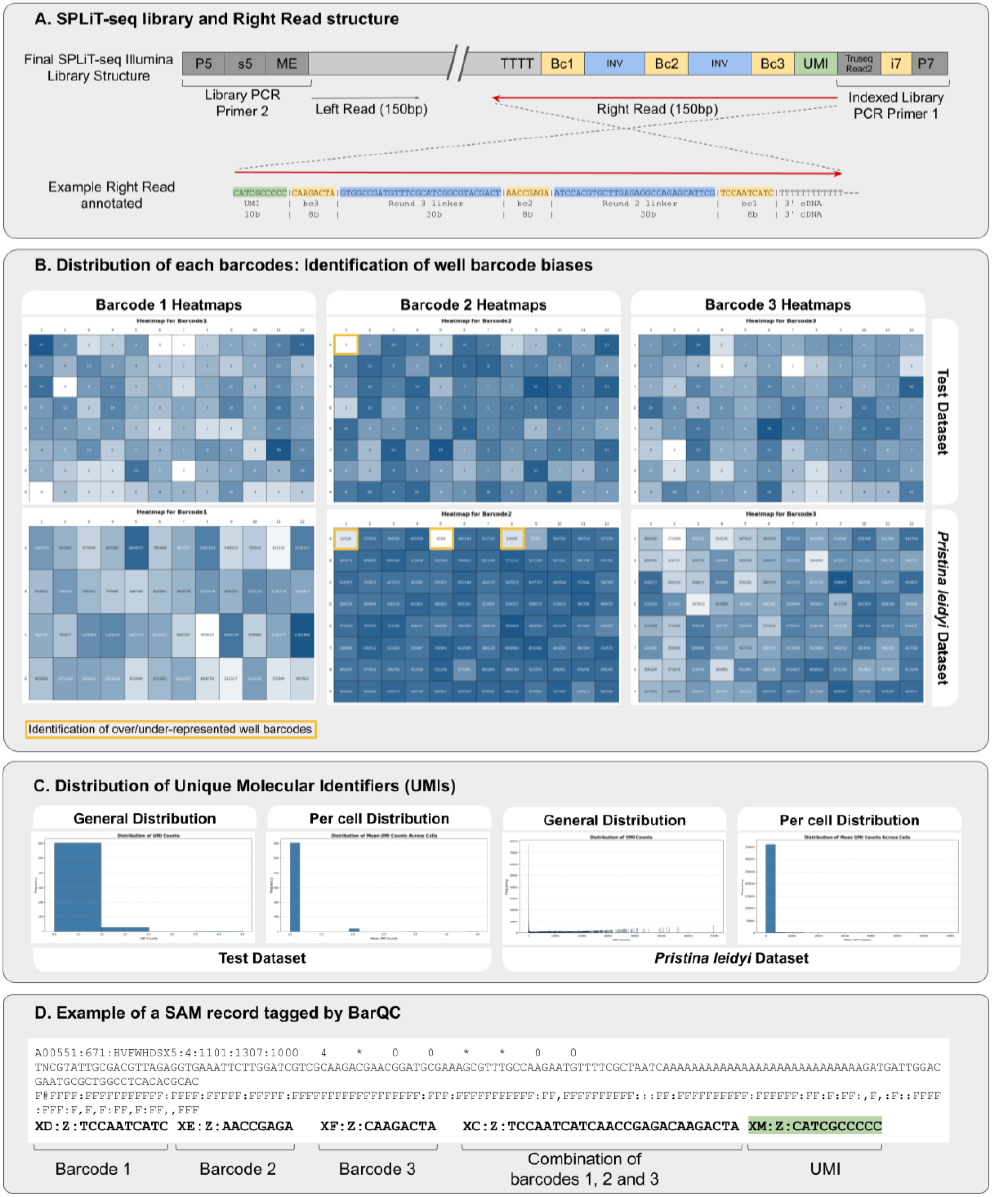
Overview of the SPLiT-seq library structure and BarQC main outputs. (A) SPLiT-seq library construction and final Right Read structure. The barcoding information is contained in the Right Reads (in 3’-5’ direction). The right reads are linked to their corresponding Left Reads (in 5’-3’ direction) which contain exclusively a part of the transcript and serve to quantify gene expression. The Right reads are composed of the UMI sequence (10bases), the barcodes 1 (10 bases), 2 and 3 (8 bases each) and finally the 3’ part of the mRNA. The barcodes are separated by two invariable regions, called round2 and round3 linkers. (B) example Heatmap outputs for barcode 1, 2 and 3 (C) Unique Molecular Identifier (UMI) distribution (general and per cell distributions). UMIs are important to control for PCR-caused biases. Lower UMI counts are indicative of a lower number of duplicated reads due to PCR amplifications. (D) Example of a SAM record tagged by barQC. The tags “XD:Z”, “XE:Z”, “XF:Z” contain the sequence of barcodes 1, 2 and 3, respectively. The tag “XC:Z” is the combination of the three barcodes and is directly associated to cellular identity. The tag “XM:Z” is the UMI sequence associated with the record. Shared UMI sequence among several records is indicative of PCR amplification inherent to the SPLiT-seq library construction and is not a biological copy of the transcript connected to the read.

Here, we introduce BarQC, a tool designed to provide comprehensive quality control (QC) metrics, including statistics on the number of barcoded reads (cell informative reads), heatmaps to visualise the abundance of each barcode across wells and plots of the Unique Molecular Identifiers (UMIs) distribution, an important metric to control for technical *versus* biological read duplicates. In addition to Quality Control, BarQC can preprocess the FASTQ reads from SPLiT-seq (by retrieving non perfect but correctable barcodes) outputting a BAM file of tagged reads.

## Implementation and outputs

BarQC is implemented using the Python3 language and incorporates memory (--memory_limit) and thread processing (-t) controls, which can be tuned by the user. Beside these options, BarQC requires as input the trimmed Left (-f1) and Right reads (-f2), an output prefix (-o) and the path to the directory containing the expected barcodes (three files, one per barcode round). BarQC is provided with a toy test dataset (976 paired reads) and the currently used set of barcode-well associations for all 3 rounds of split/pooling. However, in the case users have a different SPLiT-seq set-up, custom expected barcode files can be built, following the Comma Separated Value format (Well Position, Name, Barcode). A Conda recipe is provided for handling dependencies.

We assume proper read trimming has been performed prior to using BarQC. Although data trimming is beyond the scope of this application and left to the user’s discretion, it is essential to ensure a minimum Right Read length. The UMI and barcode region are typically 96 total bases (Figure 1A), so we recommend setting a minimum length of 96 bases for the Right Reads.

BarQC is fundamentally a Quality Control tool that outputs heatmaps of the number of reads tagged with each barcode (Figure 1B) allowing the user to have an overarching view of each barcode distribution at each “pooling” step of the SPLiT-seq protocol. This is particularly useful to detect wells with a biased number of barcodes (either associated with too many or too few reads). This information can later be accounted for in the analysis and interpretation of the data. BarQC also outputs the number of reads successfully tagged for the three barcodes, the number of complete UMIs retrieved and the distribution of UMIs (Figure 1C), which can help users understand the composition of their sequencing data. Besides providing an overview of the SPLiT-seq data, BarQC can be also run in a data processing mode, which results in a sorted BAM file containing the left read sequences, along with their respective barcode and UMI tags, as well as their combination (Figure 1D). This sorted BAM file can then be used, in conjunction with an alignment file, to build a matrix of expression. A suggested pipeline from raw SPLiT-seq reads to a raw expression matrix is proposed and depicted in Figure S3.

## Method Overview

Following initial trimming of the raw sequencing (library adaptor removal and low-quality end trimming), a prerequisite to perform downstream single cell RNA-seq analyses is to associate reads to their corresponding cells. BarQC is designed to locate and extract the sequence of the different barcodes in the Right Reads to correctly assign Left Reads to their respective cells. Our method relies on aligning the reads to the invariant sequences (30x2 bases), which link the barcodes (Figure 1A). This idea is similar to the one implemented in SCSit (Luan *et al*., 2021) and LR-splitpipe (Rebboah *et al*., 2021) However, SCSit encounters an error during execution (personal observation and (Kuijpers *et al*., 2024) and LR-splitpipe is tailored to Long read Split-seq. BarQC uses the aligner BBmap (https://sourceforge.net/projects/bbmap/, version 39.01 tested) to map the first 100 bases of the Right Read to the invariant sequences. If the read is shorter than 100 bases (e.g. due to trimming), the entire sequence is used. BBmap was chosen for its ease of use, sensitivity and very complete CIGAR string output which summarizes each alignment. In particular, BBmap allows us to set a short k-mer size (BarQC uses a default of 8) and a low minimum alignment identity (set at 0.1), which increase the sensitivity of the mapping. The CIGAR string is then parsed to find the starting position of each barcode in the read (starting indexes) (Figure S2). Based on these starting indexes, BarQC extracts each barcode sequence and compares it to the database of expected barcode sequences. If the barcode presents errors, BarQC attempts to correct them (within a hamming distance of 2) (Figure S1). Statistics of the abundance of each barcode (Figure 1B), UMI (Figure 1C) and barcode combinations are kept providing an overview of the data. If the user wishes to follow a preprocessing pipeline using BarQC, the corrected barcode sequences and UMIs are added as SAM tags to the corresponding Left Read record, in a non-aligned BAM file (Figure 1D and Supplemental Figure S1).

## Application

To showcase BarQC’s features, we tested it on a recent publicly available scRNA-seq dataset using SPLiT-seq from *Pristina leidyi* (Álvarez-Campos et al., 2024) and a test dataset shipped with BarQC. The program was run on a Linux server allowing 600G of memory and 50 threads. For the *P. leidyi* dataset (SRR24298735), raw reads were downloaded and adaptors removed with cutadapt (version 4.2) (Martin, 2011), keeping a minimum Right Read length of 96bp (complete barcode sequence length). The resulting 60,507,040 SPLiT-seq clean (trimmed) paired-reads were processed by BarQC (Figure 1), which successfully tagged 47,257,570 reads (78.10%) with all three barcodes, providing read-to-cell relationships. For the test dataset available with BarQC, 66.50% (649 of the 976 cleaned paired-reads) were successfully tagged (Figure 1). With these datasets, several biases among the different wells could be detected indicating an uneven representation of each barcode (highlighted in yellow in Figure 1B). This is particularly useful in the case of Barcode 1, as these barcodes are usually associated with biological samples or conditions. An unbalance of cell number between distinct samples is important to take into consideration in downstream analyses to avoid over or underrepresenting some samples compared to others, which could lead to misinterpretations.

Overall, BarQC is a powerful tool for analysing SPLiT-seq data, offering an overview of the dataset, including barcode and UMI distributions, while providing visual means to identify potential biases in barcode composition. Beyond QC analysis, BarQC enables pre-processing of SPLit-seq data by associating each barcode, barcode combination and UMI to their respective read. This step is essential for linking reads to their corresponding cell. Finally, by leveraging the invariant linker region positions, BarQC can locate each barcode with greater flexibility and sensitivity than methods relying on the theoretical fixed positions. This allows the potential for retrieving more informative reads.

## Acknowledgments

We are thankful to the Almudi lab for their contributions and discussions. Our thanks also go to the Solana lab for always being open to collaborative dialogue. We also thank Oriol Martínez for his help in improving data visualisation.

## Supplementary Figures

**Figure S1.**
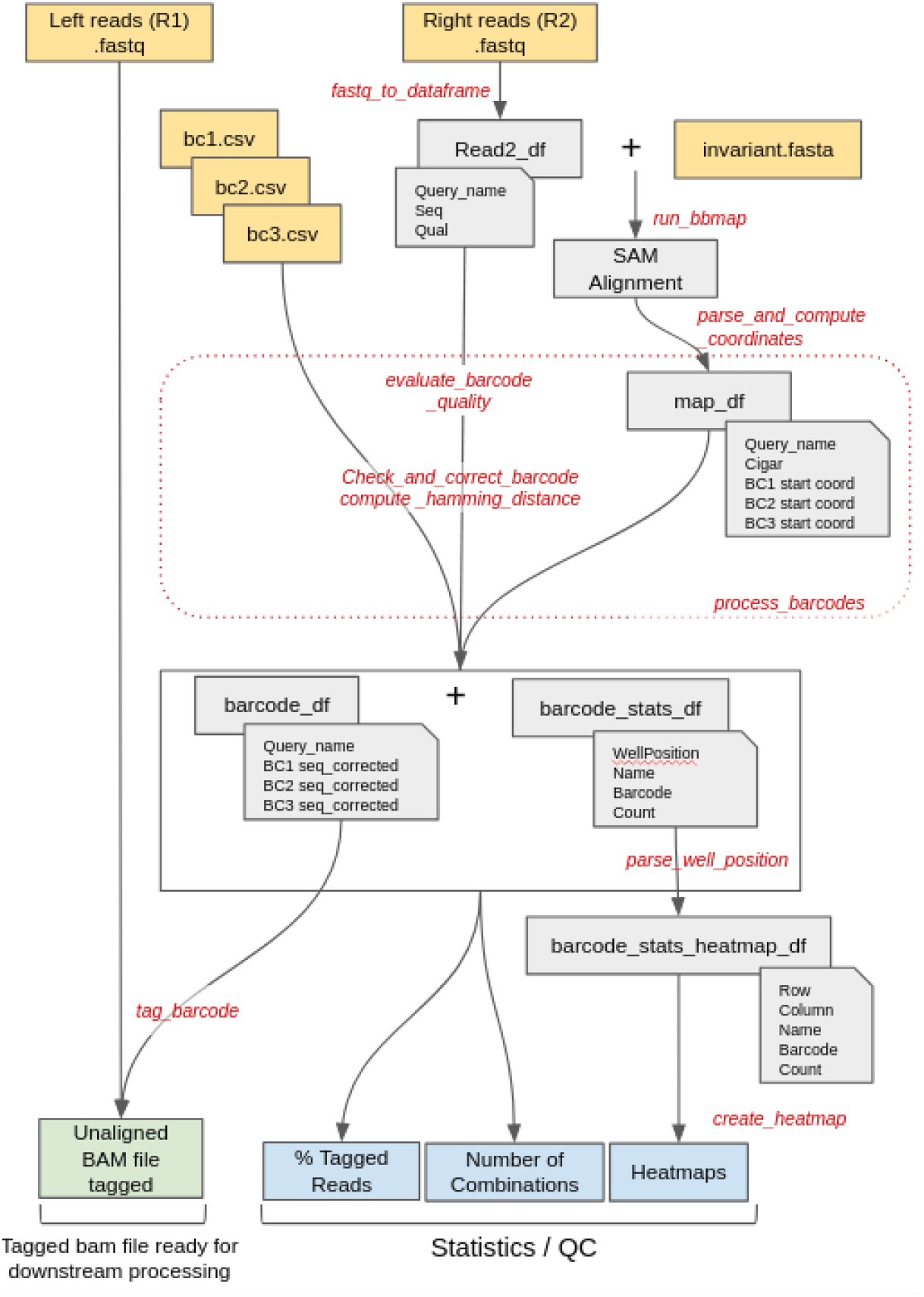
Internal procedure of BarQC. Given single-cell paired reads obtained by the SPLiT-seq method, as well as the barcodes used to tag the transcripts of cells and the invariant linker region, BarQC locates, identifies, and corrects the barcodes in the Right Reads and provides basic statistics on the number each barcode appears and for each barcode combination. It also outputs an unaligned BAM file of the Left Reads tagged for each barcode, their combination and the UMI.

**Figure S2.**
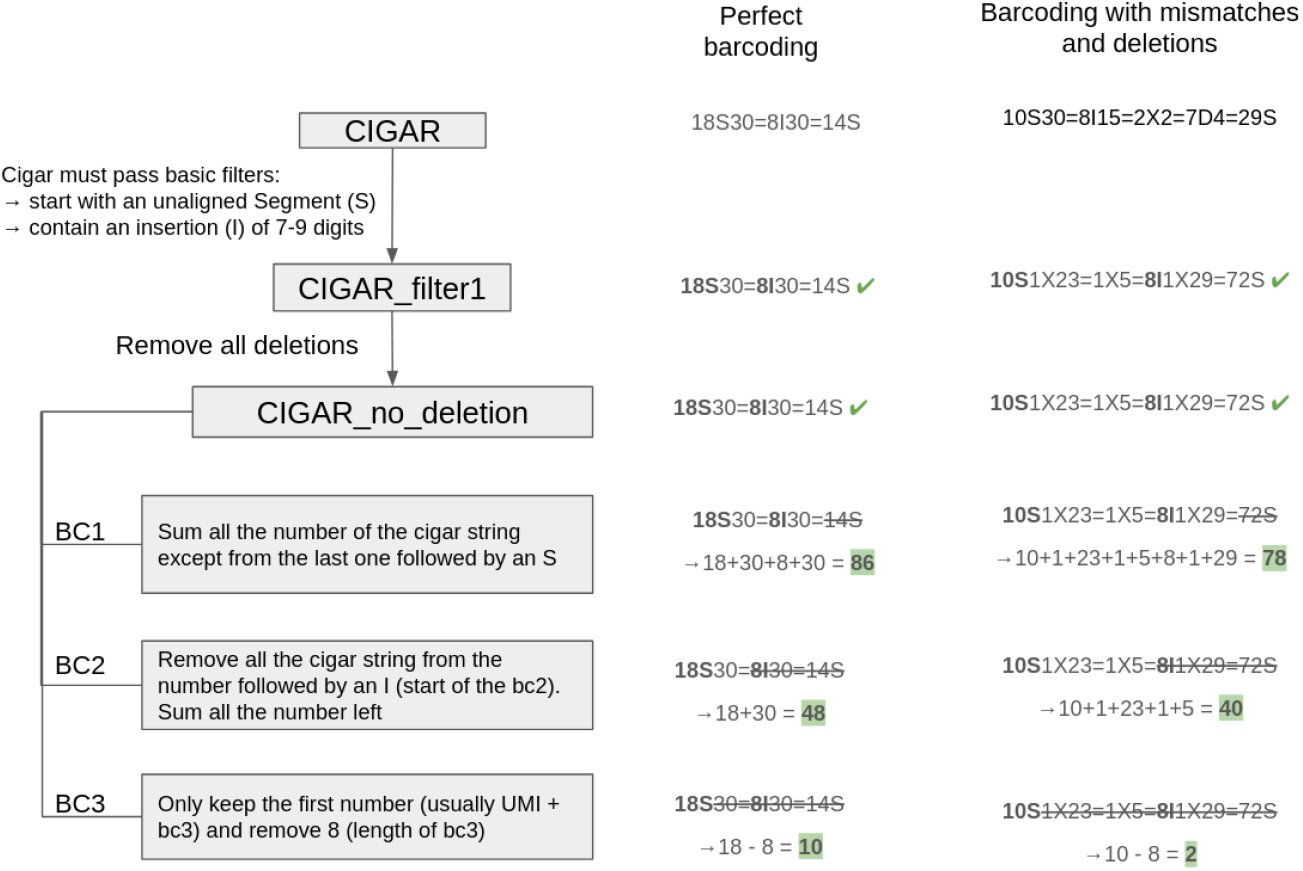
Diagram explaining the CIGAR string parsing to obtain the positions of each one of the barcodes.

**Figure S3.**
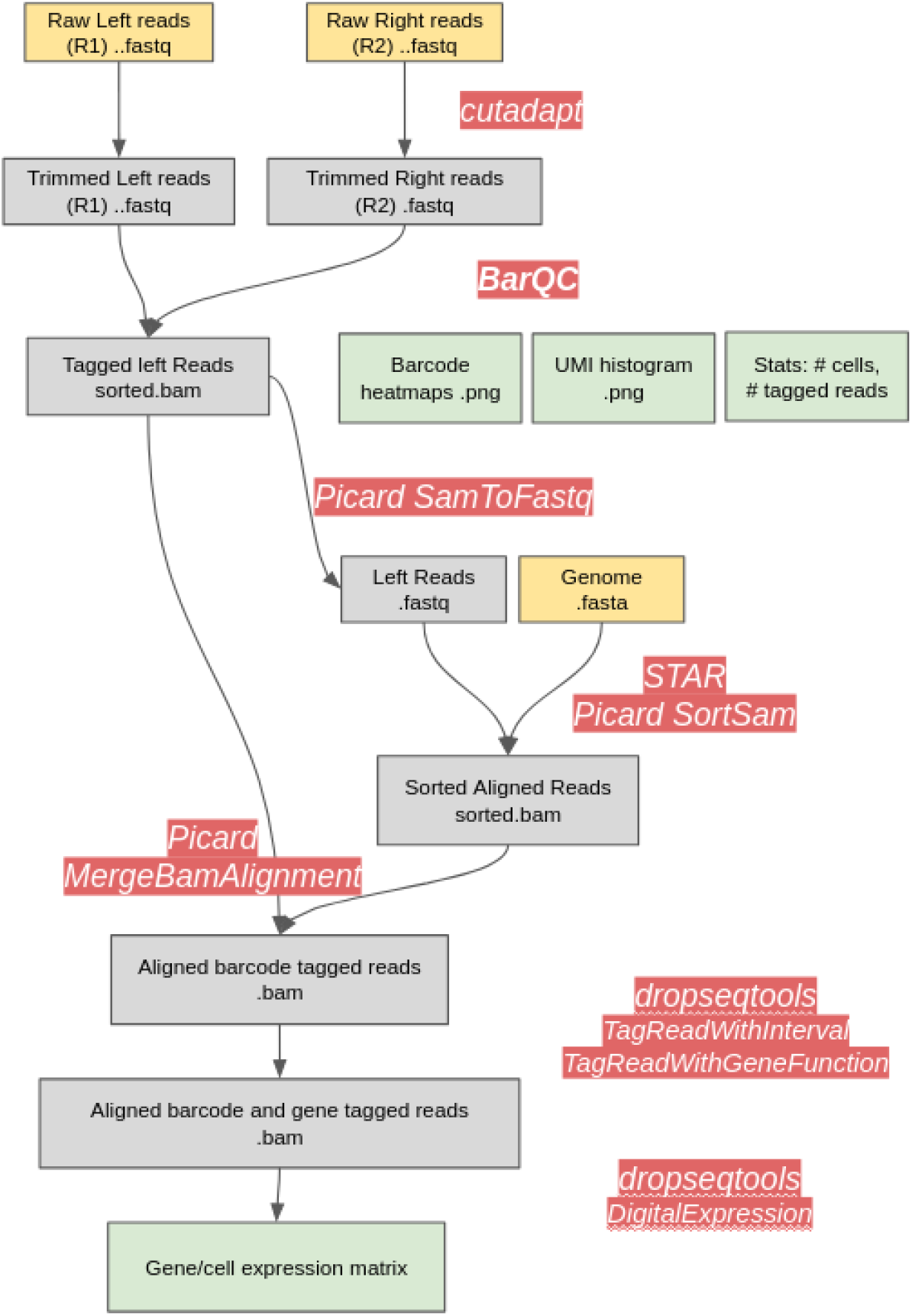
Suggested pipeline to preprocess paired-end reads generated from the SPLiT-seq method incorporating the program BarQC. This pipeline uses the dropSeqTools toolkit originally developed for single-cell sequencing based on the droplet single cell isolation method. Software is highlighted in red. Input/output files are in boxes. User inputs, Intermediate and output files are in yellow, grey, and green, respectively.

